# Maximizing Efficiency in SedaDNA Analysis: A Novel Extract Pooling Approach

**DOI:** 10.1101/2023.10.17.562718

**Authors:** Victoria Oberreiter, Pere Gelabert, Florian Brück, Stefan Franz, Evelyn Zelger, Sophie Szedlacsek, Olivia Cheronet, Fernanda Tenorio Cano, Florian Exler, Brina Zagorc, Ivor Karavanić, Marko Banda, Boris Gasparyan, Lawrence Guy Straus, Manuel R. Gonzalez Morales, John Kappelman, Mareike Stahlschmidt, Thomas Rattei, Stephan M. Kraemer, Susanna Sawyer, Ron Pinhasi

**Affiliations:** Department of Evolutionary Anthropology, University of Vienna, Austria; Human Evolution and Archaeological Sciences (HEAS), University of Vienna, Austria; Departament de Biologia Animal, de Biologia Vegetal i d’Ecologia, Universitat Autònoma de Barcelona, Bellaterra, Spain; Department of Archaeology, Faculty of Humanities and Social Sciences, University of Zagreb, Croatia; Institute of Archaeology and Ethnography, National Academy of Sciences of the Republic of Armenia; Department of Anthropology, University of New Mexico, Albuquerque, USA; Instituto Internacional de Investigaciones Prehistoricas de Cantabria at the Universidad de Cantabria; Department of Anthropology and Department of Earth and Planetary Sciences, The University of Texas, Austin, TX, USA; Division of Computational Systems Biology, Centre for Microbiology and Environmental Systems Science, University of Vienna, Austria; Department of Environmental Geosciences, Centre for Microbiology and Environmental Systems Science, University of Vienna, Austria; Institut für Analytische Chemie, University of Vienna, Austria; Forschungsverbund Umwelt und Klima, University of Vienna, Austria

**Keywords:** sedaDNA, ancient DNA, sediments, post-extraction pooling, hybridization capture

## Abstract

In recent years, the field of ancient DNA (aDNA) has taken a new direction toward studying human population dynamics through sedimentary DNA (sedaDNA), enabling the study of past ecosystems. However, the screening of numerous sediment samples from archaeological sites remains a time-consuming and costly endeavor, particularly when targeting hominin DNA. Here, we present a novel high-throughput method that facilitates the fast and efficient analysis of sediment samples by applying a pooled testing method. Our approach involves combining multiple extracts, allowing users to parallelize laboratory procedures early in the sample preparation pipeline while effectively screening for the presence of aDNA. Pooled samples that exhibit aDNA signals can then undergo detailed analysis, while empty pools are discarded. We have successfully applied our extract pooling method to various sediment samples from Middle and Upper Paleolithic sites in Europe, Asia, and Africa. Notably, our results reveal that an aDNA signal remains discernible even when pooled with four negative samples. We also demonstrate that the DNA yield of double-stranded libraries increases significantly when reducing the extract input, potentially mitigating the effects of inhibition. By embracing this innovative approach, researchers can analyze large numbers of sediment samples for aDNA preservation, achieving significant cost reductions of up to 70% and reducing hands-on laboratory time to one-fifth.

## Introduction

Genomic material can be obtained from various environmental sources, including soil (Allen et al., 2023; Ariza et al., 2023), water (Antich et al., 2021; Blattner et al., 2021), or air (M. D. Johnson et al., 2021; Lynggaard et al., 2022). Hence, environmental DNA (eDNA) is a valuable tool for ecological surveys of species that are challenging to observe directly or in the absence of their remains. The initial applications of eDNA focused on retrieving microbial genetic material from environmental samples rather than culturing them prior to extraction (Ogram et al., 1987; Olsen et al., 1986; Pace et al., 1986). Since then, numerous studies have been published, focusing on microbial taxa recovered from sediments and permafrost (Coolen & Overmann, 1998; Fierer & Jackson, 2006; S. S. Johnson et al., 2007; Shi et al., 1997; Willerslev et al., 2004). These studies have exemplified that the retrieval of DNA from archaeological sediments and permafrost could be used to study our past (Hofreiter et al., 2003; Willerslev et al., 2003). In its early stages, the analysis of ancient sediments focused on simple presence/absence studies through metabarcoding quantification (Taberlet et al., 2012; Valentini et al., 2009) involving the use of taxon-specific primers to amplify short gene regions, followed by high throughput sequencing (HTS) (Glenn, 2011). This approach has demonstrated great potential for the reconstruction of past ecosystems and the impacts of climate change (Rijal et al., 2021) from various sources, including sediments (Jørgensen et al., 2012; ter Schure et al., 2022), lake sediments (Giguet-Covex et al., 2014; Han et al., 2023; Jia et al., 2022; Rijal et al., 2021; Seeber et al., 2022), and frozen soils (Bellemain et al., 2013), even in the absence of macrofossils. Metabarcoding offers a cost-effective option. However, it does not allow further analyses such as phylogenies or population genetic tests. Alternatively, shotgun sequencing of sedaDNA provides a greater taxonomic resolution, as it analyzes all the genetic material contained within a sample, potentially allowing for the retrieval of full genomes (Gelabert et al., 2021; Parducci et al., 2019; Stahlschmidt et al., 2019). As shotgun sequencing is usually ineffective in terms of endogenous DNA sequencing, studies have applied targeted hybridization capture, particularly focusing on mammalian and hominin mitochondrial DNA (Slon et al., 2017; Zavala et al., 2021; Zhang et al., 2020) and nuclear DNA in archaeological sediments (Vernot et al., 2021). These capture approaches, however, add significantly to the expense and time needed, particularly when targeting rare hominin sedaDNA from archaeological sediments.

Even with the addition of targeted hybridization capture, sediment samples may not have any aDNA preservation. The preservation of sedaDNA in sediment is variable and depends on climatic conditions and the chemical composition of the sediment (Nagler et al., 2018; Parducci et al., 2017; Pietramellara et al., 2009; Slon et al., 2017; Zavala et al., 2021). In Denisova Cave, for example, the majority of samples exhibited the presence of faunal and hominin mitochondrial (mt)DNA (Zavala et al., 2021), while, in contrast, in Sefunim Cave, only four out of 33 tested samples showed traces of mammalian mtDNA (Slon et al., 2022). Massilani et al. were able to extract ancient mammalian mtDNA from approximately 50% of the analyzed, resin-impregnated sediment blocks (Massilani et al., 2022). This results in the potential for lost time and money when screening new sites or layers where preservation is unknown.

The potentially low success rate of sedaDNA calls for a methodological screening optimization to allow for a faster and more cost-effective assessment of samples. Sample pooling is a common practice in the field of eDNA, primarily used to assess the biodiversity of ecological communities. The concept of pooling multiple samples for collective analysis was established in the 1940s when it was initially employed to detect positive cases of Syphilis during World War II (Dorfman, 1943). Over the years, this approach has found applications in various fields, including the screening of infectious diseases such as hepatitis viruses B and C (García et al., 1996; Roth et al., 1999) human immunodeficiency virus (HIV) (Quinn et al., 2000), and, more recently, during the SARS-CoV-2 pandemic (Mutesa et al., 2021; Sawicki et al., 2021). In the context of sedimentary eDNA screening, several pooling strategies have been proposed to reduce costs, including pre-extraction (Keshri et al., 2015; Song et al., 2015; Tveit et al., 2013), post-extraction (Hestetun et al., 2021), and post-PCR pooling (Hestetun et al., 2021; Rijal et al., 2021). A recent study demonstrated the successful pooling of multiple sample libraries into a single capture reaction while optimizing reagents costs (Zavala et al., 2022). However, there is currently no experimental framework examining how post-extraction sample pooling of ancient sediments affects the behavior of the deamination signal and the abundance of authentic ancient DNA fragments.

In this study, we hypothesize that deamination signals can be maintained and detected with increased post-extraction pooling of up to five sediment samples. We further hypothesize that the abundance of unique DNA reads, mapping to our reference panel, will not significantly decrease in the various extract pools compared to the individual stand-alone sample. To test these hypotheses, we applied multiple degrees of extract-pooling to Paleolithic sediment samples from European, Asian, and African cave sites and one burial site. Our findings demonstrate that up to five extracts can be pooled while maintaining a detectable signal of aDNA and achieving an average 1.36-fold increased DNA yield across all pooling levels compared to the average unpooled state. This high-throughput pooling method will allow for large-scale sediment analyses for aDNA preservation with great reductions in costs and time.

## Results

### In silico simulation of sample pooling

Following the previous evidence for detectable sedaDNA after pooling post-PCR extracts (Rijal et al., 2021), we conducted an in-silico simulation to examine the impact of pooling on the characteristics of ancient mammalian mtDNA sequences using mitochondrial reference genomes of *Homo sapiens* and domestic goat *(Capra hircus).* To simulate the characteristics of ancient DNA we employed an online sequence manipulation tool (https://www.bioinformatics.org/sms2/split_fasta.html), fragmenting the original sequences into segments of 30, 35, and 40 base pair length. These fragmented sequences were then combined into composite files for each species, containing fragments of varying lengths. Next, we introduced deamination on the composite sequences with deamination rates set at 10, 30, and 50 percent per species. To model differential deamination levels between species within a single sediment sample, we combined human sequences with a 10 percent deamination rate with *Capra* sequences exhibiting deamination rates of 30 and 50 percent, respectively.

We carried out sequential pooling of the deaminated composite sequence files for human, *Capra,* and the combination of human and *Capra*. These pooling steps were simulated by adding random reads extracted from a sediment sample that showed no detectable aDNA reads after the initial screening process. The pooling was obtained up to a dilution of 1:10 (**Supplementary Figure S1**). We assessed deamination and read count, which are the main measures of aDNA, at each pooling step using a custom bioinformatics pipeline detailed in the methods. The simulation results demonstrated stable read numbers and deamination levels across pooling steps with a non-significant decrease in read count (p>0.05), indicating that an aDNA signal can be detected at a dilution up to 1:10-fold (**Figure 1**).

**Figure 1.**
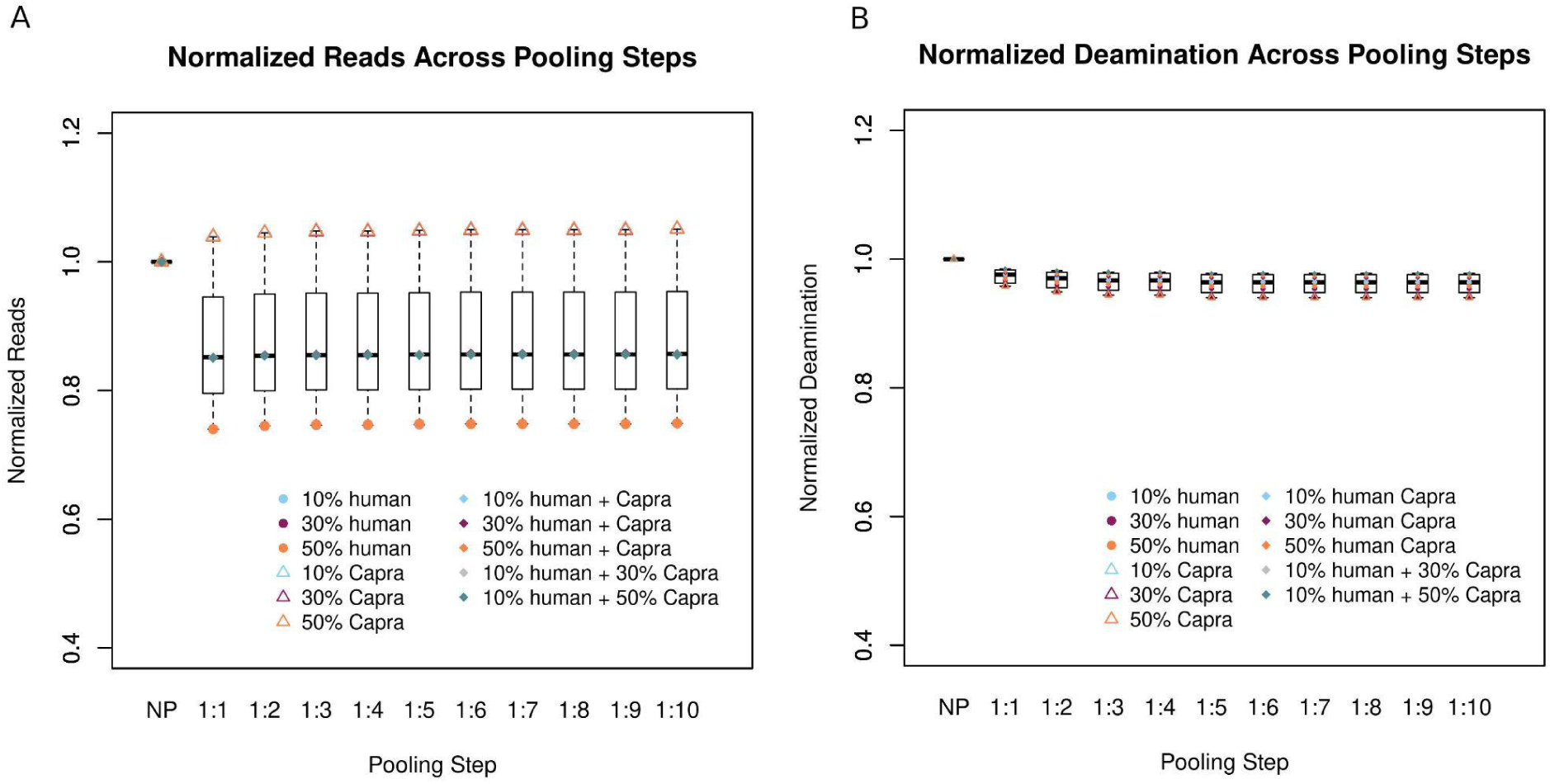
**A)** Read numbers normalized by the unpooled state for each simulated mitochondrial dataset at different pooling steps, relative to the non-pooled (“NP”) condition. **B)** Simulated deamination values normalized by the unpooled state for each simulated mitochondrial dataset at different pooling steps.

### The pooling of extracts does not decrease the sensibility

As part of the MINERVA project we screened archaeological sediments for ancient DNA employing a custom hybridization capture kit comprising 51 mammalian species (Tejero et al., 2023). Due to the age of the sequences, aDNA displays distinct signs of degradation characterized by two main molecular features: strand breaks resulting in short DNA fragments and C to T substitutions caused by deamination (Briggs et al., 2007; Brotherton et al., 2007; Sawyer et al., 2012). We classified our samples using two parameters: A) the quantity of DNA expressed as the number of unique reads and B) the damage of these reads, measured by the occurrence of C to T transitions. Based on our experience, we define four states into which sediment samples commonly fall: 1) reads with a deamination level above 0.1, and more than 1000 recovered unique reads, 2) reads with a deamination level above 0.1, and fewer than 1000 recovered unique reads, 3) reads with a deamination level below 0.1 and more than 1000 recovered unique reads, and 4) reads with a deamination level below 0.1, and fewer than 1000 recovered unique reads (**Table 1**). For convenience, all states are from here on referred to by their corresponding reference number (1-4). We further defined state 1 as the threshold for a positive sediment sample (i.e., deamination level above 0.1 and 1000 unique endogenous reads, aligning to the mammalian reference genome panel used during the capture process). A failed sample is defined as a sample falling into states 2-4. In a different study on sedaDNA, a sample was considered to be positive for ancient hominin DNA if the number of DNA fragments assigned to hominins constituted at least 1% of the total identified fragments and if it had at least 10 fragments showing deamination which was significantly higher than 10 % (Zavala et al., 2021). Another study also used a 10% deamination frequency at both ends as a cut-off (binomial CI ≥ 10%) (Vernot et al., 2021).

**Table 1:**
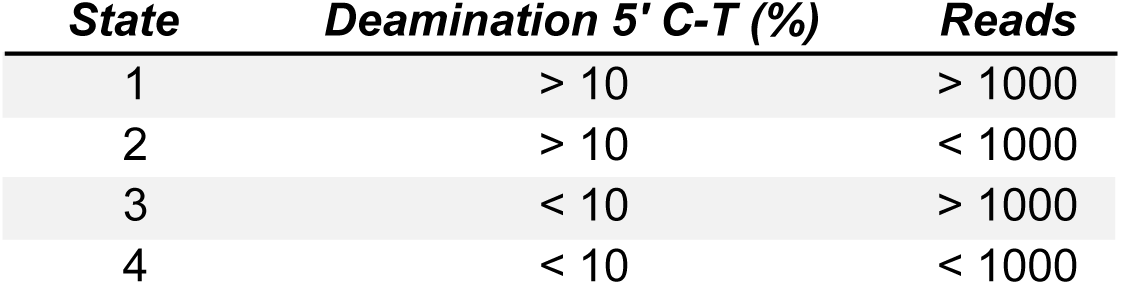
The summary of the four states in which we classify the positive samples.

| **State** | **Deamination 5' C-T (%)** | **Reads** |
| --- | --- | --- |
| 1 | > 10 | > 1000 |
| 2 | > 10 | < 1000 |
| 3 | < 10 | > 1000 |
| 4 | < 10 | < 1000 |

To evaluate our pooling simulation in a laboratory setting, we selected eight test samples to represent these four distinct states (two samples per state according to earlier screening analyses). However, two of the test samples had to be excluded from the study. One test sample falling into state 2 was contaminated with 1.67x modern human DNA. Additionally, the screening results for a state 3 sample could not be reproduced; the sample displayed higher deamination after re-extraction. A negative sample was chosen for the pooling. This sample had an average deamination of 4.7% at both ends and 651 reads aligning to our mammalian mtDNA panel. The extract of each test sample was pooled with the extract of our chosen negative sediment sample in increasing amounts up to a dilution of 1:4 (**Figure 2**). This process was performed in two sets for each state, with two distinct candidate samples meeting states 1-4. To ensure accuracy, each dilution level was replicated three times.

**Figure 2.**
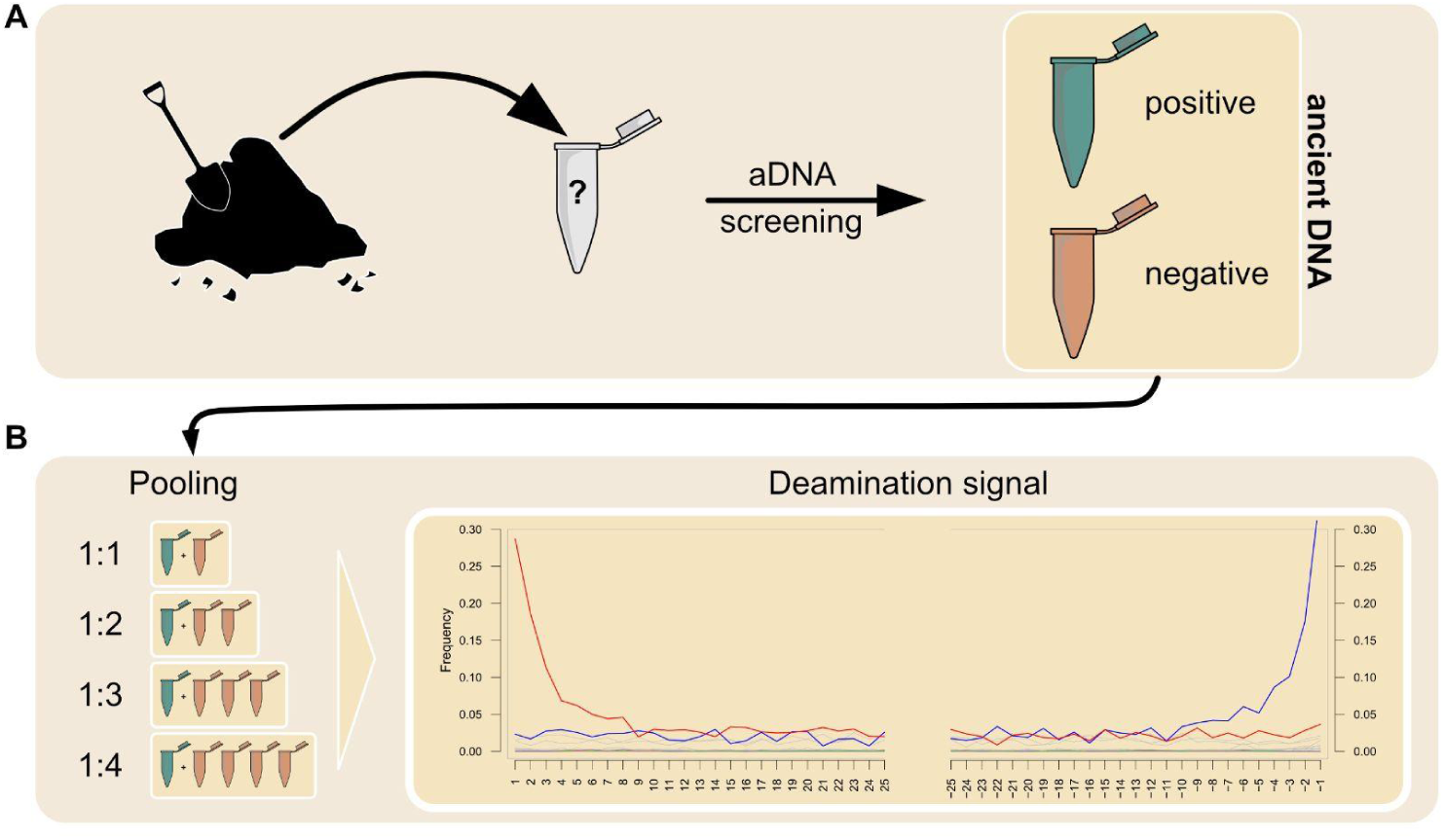
Sediment samples were individually screened to distinguish samples with preserved ancient DNA (positive) from samples devoid of aDNA (negative). The extracts of positive samples were pooled with up to 4 extracts of a negative sample to recreate a pool testing method. The presence of an aDNA signal was evaluated by the deamination signal and the number of unique reads.

The experiment showed that for states 1 (R^2^ = – 0.73), 3 (R^2^ = 0.45), and 4 (R^2^ = – 0.86) deamination remains stable with increased pooling (**Figure 3B**). Even at low deamination and low read levels (state 4), the signal for aDNA could be maintained. State 2 (R^2^ = – 0.94) showed a significant decrease in deamination level with increased pooling (p = 0.01789).

**Table 2:** Comparison of DNA yields in normalized, average read numbers across pooling levels.

| ***Pooling level*** | ***avg reads*** | ***% increase in reads*** | ***fold increase in reads*** |
| --- | --- | --- | --- |
| NP | 542 | N/A | N/A |
| 1:1 | 634 | 17 | 1.17 |
| 1:2 | 798 | 47 | 1.47 |

| *Pooling level* | *avg reads* | *% increase in reads* | *fold increase in reads* |
| --- | --- | --- | --- |
| NP | 542 | N/A | N/A |
| 1:3 | 722 | 33 | 1.33 |
| 1:4 | 786 | 45 | 1.45 |
| avg of all pooling levels | 735 | 36 | 1.36 |

We observed an increase in normalized average read numbers when pooling was implemented (**Figure 3A**). Despite considerable variation within each pooling level (SD = 0.12 – 0.23), there was a significant increase in average reads between the non-pooled state and the 1:3 dilution level (*p =* 0.018) as well as the 1:4 dilution level (*p =* 0.011). DNA yields were on average increased by 17, 47, 33, and 45 percent for pooling levels 1 to 4, respectively. Overall, the DNA yield across all pooling levels was increased by 1.36-fold, compared to the average unpooled sample.

**Figure 3:**
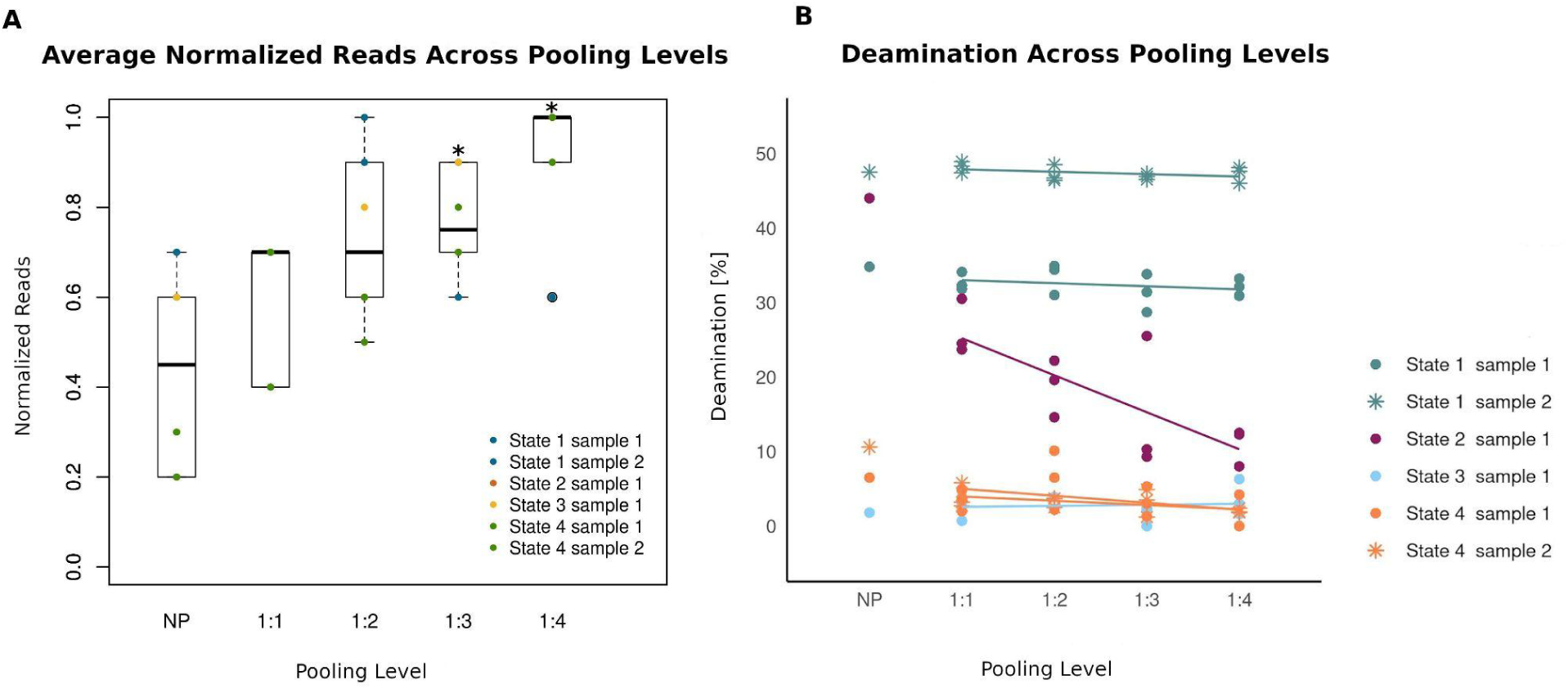

### A cost-efficient method

The screening of sedaDNA samples is a laborious and expensive undertaking. We have modeled multiple situations in order to gain insights into the cost and labor reduction of this approach. We estimate that under conditions where 10% of the samples exhibit the characteristics previously defined as positive, the implementation of our approach would result in a 50% reduction in the number of libraries prepared. In the case of a 5% positivity rate, this would amount to a 75% decrease in the number of prepared libraries. Thus, we suggest that the pooling of five extracts is a cost-effective strategy in scenarios where the prevalence of positive samples is less than 20% among the total (**Table 3**).

**Table 3:** Summary of the cost-effectiveness of the pooling strategy.

|  | **Pooling Approach** | **Non-pooling Approach** |
| --- | --- | --- |
| **Cost implication** | Substantial cost reduction (ca. 70%) and enhanced cost efficiency. | Higher cost |
| **Speed of downstream analysis** | Accelerated downstream analyses due to parallelized lab steps (one-fifth of lab time) | Slower processing and longer turnaround times |

## Discussion

### Experimental extract-pooling reduces inhibition

With our extract-pooling approach, we set out to define a cost-efficient and high-throughput method for screening archeological sediments for aDNA. When pooling was implemented, we observed a cost reduction of approximately 50-75%, including costs for reagents and sequencing. Hands-on time in the lab was reduced to one-fifth. We demonstrate that the pre-PCR pooling of Paleolithic sediment extracts does not reduce the number of aDNA reads detected in double-stranded libraries (Meyer & Kircher, 2010) across all pooling levels: extract-pooling in fact significantly (p = 0.0001) increases the efficiency of aDNA recovery across all four states on average by 1.36-fold (**Figure 3A**). The library preparation process for pooling levels incorporated different extract input volumes: 25 μL for the unpooled state, 12.5 μL for a 1:1 pooling, 8.33 μL for a 1:2 pooling, 6.25 μL for a 1:3 pooling, and 5 μL for a 1:4 pooling state. The observed increase in unique DNA fragments while reducing the extract input volume may be attributed to substances present in the sediment exerting a negative effect on PCR amplification. Common PCR inhibitors present in the organic component of soil include plant-derived substances such as tannic, fulvic, and humic acids (Hering & Morel, 1988; Matheson et al., 2014; Sutlovic et al., 2008). Such inhibitory substances can be co-extracted with DNA and interfere with enzymatic reactions in downstream lab procedures, including quantification via real-time PCR and PCR amplification due to inhibition of polymerase activity (Uchii et al., 2019). This can lead to an increased number of false negative results (McKee et al., 2015; Opel et al., 2010). The individual identification of inhibitory agents in different types of sediment in order to adapt extraction, purification, and PCR protocols is therefore necessary. A study conducted on bones and teeth demonstrated that the preparation of single-stranded libraries with minimal extract input effectively mitigates the effects of inhibition (Glocke & Meyer, 2017). To our knowledge, inhibition in ancient sediment DNA samples has never been investigated for double-stranded libraries (Meyer & Kircher, 2010). Our results show that a reduction of extract input into double-stranded libraries reduces potential inhibition significantly when comparing the read numbers of the unpooled sample with those of pooling level 1:3 (p = 0.018) and pooling level 1:4 (p = 0.011) (**Figure 3A**). The extensive dilution of a DNA extract (often up to 400 fold), is a common practice across different scientific disciplines to circumvent inhibition (Imaizumi et al., 2005; McKee et al., 2015; Schneider et al., 2009; Shutler et al., 1999). As a consequence, the target DNA will simultaneously be diluted along with the inhibitory substances (Bustin et al., 2009; Lebuhn et al., 2004). In cases of underrepresented taxa, a dilution may even result in the failed detection of endogenous DNA (Jane et al., 2015; Lindberg et al., 2007). Another study, therefore, urges a moderate extract dilution of 40 – 60-fold to effectively reduce qPCR inhibition (Wang et al., 2017). The minimal extract volume converted into double-stranded libraries in our study was 5 µL, which equals a 1:4 dilution of the regular input. In our approach, we intentionally pooled a positive sediment extract with increasing volumes of a negative extract. However, it is important to note that when applying pooling to several previously unscreened extracts, in which case the extent of inhibitors is unknown, we recommend pooling at small volumes. Another unknown factor in unscreened extracts is the extent of DNA preservation. Therefore, we cannot make a generalized conclusion that dilution consistently leads to increased DNA yields, as it relies on the assumption that the concentration of humic substances corresponds to DNA content, which would require further testing.

Further experiments will be necessary to test the optimal volume of extract converted into libraries to examine the potential for even smaller volumes than those employed in this study for enhanced DNA recovery. We recommend conducting similar experiments to assess the optimal input concentrations of both extracts and pre-PCR libraries in the context of ancient DNA.

### Pooling vs. modern contamination

The extent of deamination remains stable as pooling increases, with one exception observed in state 2 (deamination levels above 10%, read numbers below 1000) (**Figure 3B**). For samples exhibiting the characteristics of state 2, it is plausible that the introduction of increased dilution with the negative sample containing a 2.4x modern human contamination masks the impact of the limited aDNA portion within the positive sample. This aligns with the observations in Figure 3, demonstrating a gradual reduction in C to T transitions with each increment in pooling. Notably, a parallel observation can be made for another sample categorized under state 2 which was ultimately excluded from the dataset due to a 1.67x modern human contamination. This sample displayed a substantial reduction in its deamination signal as pooling intensified (**Supplementary Figure S2**). This result supports the hypothesis that modern DNA can effectively mask the distinctive markers of aDNA: The various iterations of PCR amplification during the laboratory pipeline can introduce a technical bias towards the amplification of the more abundant and less fragmented modern DNA. Characteristic aDNA damage patterns including miscoding lesions and single-strand breaks (Lindahl, 1993) can block polymerases, leading to chimeric sequences or the termination of amplification (Pääbo et al., 1989). Consequently, this phenomenon leads to the prioritized sequencing of modern sequences over ancient ones during the screening process.

### Finding the right mix – how many samples to pool?

In this study, we employed equal-volume pooling of non-normalized extracts (according to concentration) to achieve increased pooling levels. Rather than merging different extracts, we utilized a single (positive) extract and combined it with progressively larger volumes of another (negative) extract to attain higher pooling levels. This strategy was used to simulate the potential low success rate of sedaDNA samples (Massilani et al., 2022; Slon et al., 2022). However, it’s important to note that equal-volume pooling does not guarantee equal representation of each extract within the pool. In our experimental setup, this was negligible, however, when applying our extract-pooling method for the screening of different sediment samples, equal mass pooling is not advised as it does not circumvent heterogeneity in the extract pool. To address this, we propose assessing the quantity of double-stranded DNA in individual extracts through the use of a fluorometer before pooling and combining extracts at equal concentrations to ensure a more balanced representation in the pooled libraries.

Despite several studies that have evidenced that extracts can be pooled in the order of hundreds (Rijal et al., 2021) when the objective is to identify positive samples for downstream analyses, an elevated number is suboptimal as a large (financial) effort is required to produce individual libraries from sample pools that tested positive. Therefore, it is clear that an equilibrium must be found. In our laboratory experiment, we established an upper threshold of five extracts per individual pool. This decision was reached to ensure a balance between analytical efficiency and resource optimization. However, it is worth noting that there is potential for further exploration in future studies regarding the optimal number of samples that can be pooled.

Recently, there has been another notable methodological advancement aimed at enhancing the cost-effectiveness of large-scale sample screening, focusing on a step much closer to the end of the laboratory pipeline, referred to as multiplex capture (Zavala et al., 2022). In contrast to our method, multiplex capture involves individual library preparation, double-indexing, and initial amplification. The streamlining of lab processes is achieved at the step of hybridization capture, through the pooling of multiple double-indexed libraries into a single capture reaction. While reducing the costs for capture reactions considerably, the majority of lab processes have to be conducted individually. This method could additionally be tested in combination with our extract-pooling method.

In this study, we have successfully demonstrated two key findings that have important implications for the field of sedaDNA research: 1) Preservation of sensitivity: Our study shows that the pooling of up to five extracts did not compromise the sensitivity of ancient DNA detection. This finding implies that extract pooling can be employed as a cost-effective and efficient strategy for large-scale sediment sample screening without sacrificing the ability to detect ancient DNA. 2) Effective inhibition reduction: Our results highlight that inhibition in double-stranded libraries can be significantly reduced by using low extract volumes. This observation is critical as it addresses a common challenge in ancient DNA research – the presence of inhibitory substances in sediment samples. This approach allows researchers to optimize extract input volumes, leading to enhanced recovery of ancient DNA while minimizing the impact of potential inhibition. Overall, this cost-effective and high-throughput approach has the potential to advance the field and facilitate the study of ancient ecosystems and populations.

## Materials and methods

### DNA extraction and library preparation

Sediments were obtained from multiple Paleolithic cave sites and one burial site: (El Mirón (Spain), Velika Pećina – Kličevica (Croatia), Hovk-1 (Armenia), Krems-Wachtberg (Austria), and Shinfa-Metema 1 (Ethiopia) (**Supplementary Information S1**). To mitigate the risk of modern contamination, we implemented strict measures such as the use of gloves, disposable lab suits, and other sterile equipment. All of the samples were prepared in dedicated clean room facilities at the University of Vienna. We included negative controls at each step (extractions, libraries, and PCR) to monitor potential reagent contamination. During the screening for candidate samples, amounts of 50-60 mg of sediment were used for DNA extraction following the protocol from Dabney (Dabney et al., 2013) with adaptations from Korlević (Korlević et al., 2015), and eluted in 50 µL TET buffer (10 mM Tris-HCl, 1 mM EDTA, 0.05% Tween 20, pH 8.0). Double-stranded libraries were prepared of half of the extract as described in (Meyer & Kircher, 2010), omitting the shearing of DNA into smaller fragments. Instead of SPRI beads, a MinElute PCR Purification kit from Qiagen was used for cleaning up the samples and eluted in 40 µL EBT buffer (1 mM EDTA, 0.05% Tween-20). We added a positive control using 24 µL of deionized water and 1 µL of a 1:250 dilution of CL104.

### Mammalian mtDNA hybridization capture and sequencing

The number of PCR cycles was determined individually for each library using a qPCR machine (AriaMx Agilent). We double-indexed (Kircher et al., 2012) and amplified 25% of each library with PfuTurbo Cx HotStart DNA Polymerase from Agilent. Amplification products were cleaned up using NGS clean-up magnetic beads (Macherey-Nagel), introducing a size selection through a concentration of 1.2x beads per sample and eluted in a total amount of 25 µL EBT buffer (1 mM EDTA, 0.05% Tween-20). In preparation for the subsequent capture, we further amplified 3x2 µL of each indexed library using KAPA HiFi HotStart DNA Polymerase and the primer pairs IS5/IS6 (Meyer & Kircher, 2010) at a rate of 20 cycles and cleaned up with magnetic beads. The concentration of the PCR product was measured on a Qubit.

We enriched for 51 mammalian mitogenomes, including human mitochondrial DNA, using custom-designed capture probes from TWIST (Tejero et al., 2023) following the TWIST capture protocol (Rohland et al., 2022). After a hybridization time of 16 hours at 65°C and several rounds of washing the enriched DNA was cooked off the probes in a PCR cycler at 95°C for 5 min (including a heated lid at 110°C). A qPCR was performed to determine the correct number of PCR cycles for each library. Half of the captured library was re-amplified using KAPA HiFi HotStart DNA Polymerase and the primer pairs IS5/IS6 and cleaned up with magnetic beads. Library concentration (ng/µL) and molarity (nmol/L) were determined through Qubit 4 and Tapestation 4150. For sequencing on a NovaSeq SP lane, libraries were pooled by including 20 ng of DNA for each and the amount for blanks was calculated to approximate 200,000 target reads per blank. The sequencing was performed at the Vienna BioCenter Core Facilities in Vienna.

### Pooling experiment

To generate sufficient extracts for the pooling experiment, we employed a modified extraction protocol that involved processing 1 g of sediment for the negative samples (i.e., those without detectable aDNA) and 250 mg of test samples (i.e., those with detectable aDNA). Following the protocol described in Dabney et al. (2013) and Korlević et al. (2015), we eluted the extracts in TET buffer, using 1 mL and 250 mL for negative and test samples, respectively. To ensure reproducibility of results across all samples, we conducted downstream lab analyses using the same methods as previously described, and carried out shallow sequencing on a NovaSeq SP lane, alongside other samples. Any sample for which the screening results were reproducible was included in the experimental set-up. In cases where the screening results could not be replicated, samples were substituted with new candidate samples. The extracts of the previously sequenced sediment samples which showed proof of aDNA were diluted with the extract of a negative sample in increasing amounts (ranging from 1:1 to 1:4) up to a total volume of 25 µL. Each pool of extracts was prepared in three replicates and converted into double-stranded libraries according to (Meyer & Kircher, 2010). We proceeded with the downstream analyses (PCR, mammalian capture, and quality controls) as described above.

### Bioinformatic processing

First, we removed the adapter sequences using Cutadapt 4.0 (Martin, 2011). The clipped reads were mapped to a composite reference file encompassing the 51 mammalian mt genomes included in the hybridization capture (Tejero et al., 2023) using bwa aln version 0.7.17-r1188 (Li & Durbin, 2010) with an edit distance of 0.01, a gap penalty opening of 2 and seeding disabled. Reads were filtered using samtools (Li et al., 2009) retaining the ones with average mapping qualities above 30. We removed duplicate sequences with samtools rmdup and assessed damage levels through mapdamage 2-2.2.1 (Jónsson et al., 2013). The quantity of the accepted reads was examined using Qualimap v.2.3 (Okonechnikov et al., 2016).

### Statistical analyses

All statistical analyses were performed in Rstudio Version 1.1.414 (Team, 2015) using R version 3.6.3 (R Core Team, 2013). All plots were produced using R Studio version 3.6.3 (2020-02-29). Boxplots were produced using the default settings: The boxes represent the Interquartile Range (IQR) and span from the first quartile (25th percentile) to the third quartile (75th percentile) of the data. The midline of the box is the median. The whiskers extend up to 1.5 times the IQR from both ends of the box to the furthest datum within that distance. Data plotting beyond that distance are represented as individual points and considered outliers. P-values were calculated in R studio using a paired Wilcoxon rank sum test with continuity correction and considered to be significant at a p-value below 0.05.

## COMPETING INTEREST STATEMENT

The authors declare that they have no competing interests to disclose.

## ACKNOWLEDGMENTS

This study was supported by a grant from the internal Research Platform MINERVA (https://minerva.univie.ac.at). P.G. was funded through COST iNEAL STSM Grant CA19141-8d068698. S.K.S. was funded by the Austrian Science Fund (FWF) M3108-G. The excavation and sampling of Velika Pećina in Kličevica was financially supported by the Croatian Science Foundation (NECEM, HRZZ-IP-2019-04-6649) and M.S. received funding from the European Research Council (ERC) MicroStratDNA project under Grant Agreement No. 101042570.

The authors would like to express their gratitude to the following individuals and institutions for their invaluable contributions to this research: Paul Knabl for the artistic expertise demonstrated in creating the figures. Natalija Čondić and Archaeological Museum Zadar for various forms of assistance during the sampling of Velika Pećina in Kličevica.

## Author Contributions

Study conceptualization: S.K.S., P.G., V.O., R.P. Methodological design: V.O., S.K.S., P.G., R.P., O.C., Experimental work: V.O., F.B., S.F., E.Z., S.Sz., F.TC., F.E., B.Z. Data analysis: V.O., P.G., and S.K.S. Archaeological conceptualization and sampling: I.K., M.B., L.G.S, MR.GM., B.G., M.B., JW.K. Project administration and funding acquisition: R.P., T.R., S.M.K., M.S. Writing: V.O., S.K.S., P.G., R.P. with input from the rest of the authors.

## Notes

### Competing Interest Statement

The authors have declared no competing interest.

